# *Ehrlichia ruminantium* infection is associated with tissue-specific microbial community shifts in *Amblyomma gemma* ticks from cattle in Kenya

**DOI:** 10.64898/2026.05.26.727963

**Authors:** Dennis Getange, Samson Mukaratirwa, Dorcas Chebet, James Kabii, Rua Khogali, Jandouwe Villinger

## Abstract

Tick-borne pathogens can reshape vector microbiomes in ways that influence pathogen colonisation and transmission, yet the interplay between *Ehrlichia ruminantium* and the microbiota of its tick vectors remains uncharacterised. We profiled bacterial communities in haemolymph, midgut, and salivary glands of infected (n = 11) and uninfected (n = 12) *Am. gemma* ticks, a vector of *E. ruminantium* in East Africa, collected from cattle in Kajiado County, Kenya, using near-full-length 16S rRNA gene amplicon sequencing on the Oxford Nanopore platform. Community composition, alpha and beta diversity, co-occurrence networks, keystone taxa, and PICRUSt2-inferred functional profiles were compared across tissue–infection status groups. We identified 226 bacterial genera dominated by *Coxiella*, *Pseudomonas*, *Acinetobacter*, *Proteus*, and *Rickettsia*. Infection was associated with tissue-specific shifts in community composition (PERMANOVA *R*² = 0.14, *p* < 0.001) and co-occurrence network structure, with midgut networks showing complete hub taxon turnover (Jaccard = 0.000, *p* = 0.043). Haemolymph communities converged around *Luteimonas* as a keystone taxon, while opportunistic Proteobacteria, including *Acinetobacter* and *Serratia*, emerged as keystones in infected midgut. Endosymbiotic *Rickettsia* was near-absent in infected tissues (0.3% vs 9.3% mean relative abundance in midgut), consistent with competitive exclusion. Functional inference identified FDR-significant enrichment of predicted aerobactin siderophore biosynthesis, antimicrobial efflux, and oxidative stress response gene families in infected microbiota. These findings show tissue-specific restructuring of the *Am. gemma* microbiome associated with *E. ruminantium* infection and point to candidate targets for microbiome-based interventions against heartwater.

**Importance:** Heartwater, caused by the bacterium *Ehrlichia ruminantium* and transmitted by *Amblyomma* ticks, kills up to 90% of susceptible ruminants and is one of the most devastating tick-borne diseases in sub-Saharan Africa. Controlling heartwater requires understanding how the pathogen interacts with the microbial communities living inside its tick vector. In this exploratory study, we show that *E. ruminantium* infection is associated with tissue-specific shifts in the *Amblyomma* tick microbiome, including reduced abundance of beneficial symbionts, elevated representation of opportunistic bacteria among community hubs, and enrichment of iron acquisition and antimicrobial resistance functions. The midgut, the first tissue colonised during infection, showed the most marked structural reorganisation. These tissue-resolved microbiome signatures point to potential targets for novel control strategies, such as anti-microbiota vaccines or approaches that reinforce natural colonisation resistance, offering new strategies to reduce heartwater transmission and protect livestock livelihoods across Africa.

## INTRODUCTION

Tick-borne diseases (TBDs) represent one of the most significant threats to global livestock production, causing billions of dollars in economic losses annually through direct mortality, reduced productivity, and control costs (1, 2). Among these diseases, heartwater caused by the obligate intracellular bacterium *Ehrlichia ruminantium* (order Rickettsiales) stands as one of the most economically devastating tick-borne diseases across sub-Saharan Africa, Indian Ocean islands and the Caribbean (3–5), affecting domestic and wild ruminants with case fatality rates reaching 90% in naive animals (6, 7). This pathogen is transmitted primarily by *Amblyomma* ticks (e.g., *Amblyomma gemma*, *Amblyomma hebraeum*, *Amblyomma variegatum, Amblyomma lepidum*), with recent evidence suggesting potential transmission by *Rhipicephalus microplus* in West Africa (8, 9). *Ehrlichia ruminantium* has developed sophisticated adaptations to persist within its arthropod vectors, demonstrating transstadial patterns that require horizontal acquisition from infected vertebrate hosts during blood-feeding (10, 11), followed by colonisation of multiple tick tissues including the gut, salivary glands, and haemolymph. Transovarial transmission, which is characterised by a disease agent being transferred from an infected female to her eggs, may rarely occur, but this has been reported in *Amblyomma hebraeum* (12) and *Rh. microplus* (8).

In Kenya, only *Amblyomma gemma*, *Am. variegatum*, and *Am. lepidum*, which maintain the bacterium solely through transstadial transmission (from larvae to nymphs and adults, or from nymphs to adults), are known to be competent vectors for *E. ruminantium* (13). *Ehrlichia ruminantium* has been detected in Kenyan ticks across multiple surveillance studies, with at least two distinct strains co-circulating in ticks from cattle and prevalence varying by location and season (14–21). Serological evidence further supports widespread *E. ruminantium* circulation, with PC-ELISA revealing high rates and levels of seropositivity in camels from Kenya, particularly among animals sampled in areas where heartwater-like disease had been reported, while co-grazing sheep in these regions also demonstrated extremely high seropositivity rates (22). Clinical heartwater-like disease associated with *Ehrlichia* spp. closely related to *E. ruminantium* has also been implicated with camel disease in Kenya (23). In addition to *E. ruminantium*, *Amblyomma* ticks harbour complex microbial communities, including diverse *Rickettsia* species (*Rickettsia africae* and *Rickettsia* endosymbionts), *Coxiella*-like endosymbionts, and other bacterial taxa that may influence pathogen dynamics (17, 24–26). These ticks also carry non-pathogenic microorganisms, which include commensal microbes acquired from the environment, as well as transovarially-transmitted endosymbionts (27–29), collectively referred to as the tick microbiota. Understanding these pathogen-microbiome interactions is a critical knowledge gap for heartwater epidemiology and control in Kenya.

Tick microbiome represents an important mediator of pathogen colonisation and transmission, as newly acquired pathogens must navigate hostile environments dominated by established microbial communities and tick immune responses (30, 31). Recent studies have demonstrated that tick-borne pathogens (TBPs) actively modulate microbiome composition of their vectors to create favourable conditions for colonisation, as observed with *Borrelia burgdorferi* in *Ixodes scapularis*, *Rickettsia helvetica* in *Ix. ricinus* and *Anaplasma marginale* in *Rh. Microplus* salivary glands and ovaries (32–34). Recent network analyses further reveal that pathogen infection can compromise colonisation resistance by reducing microbial network robustness, as demonstrated with *Anaplasma phagocytophilum* in *Ix. scapularis*, where infected ticks showed increased vulnerability to additional microbial invasions and potential co-infections (35). Rickettsial exclusion phenomena documented in other tick species suggest that endosymbiotic *Rickettsia* can block transmission of a pathogenic *Rickettsia* through transovarial interference (36, 37). However, the interactions between *E. ruminantium* and co-occurring bacteria in *Am. gemma* remain poorly understood, potentially involving direct competition for cellular niches or facilitative relationships that enhance pathogen survival. Understanding these microbe-pathogen interactions has important implications on tick control, with emerging research exploring anti-microbiota vaccination strategies targeting tick endosymbionts to reduce vector competence and block pathogen transmission (33).

This study investigated whether *E. ruminantium* infection modulates the bacterial microbiome composition and functional potential within different tissues of *Am. gemma* ticks collected from cattle in southern Kenya. Using tissue-specific 16S rRNA gene amplicon sequencing of haemolymph, midguts, and salivary glands from both infected and uninfected ticks, we: (1) characterised microbial composition and diversity within haemolymph, midgut and salivary glands of *Am. gemma* ticks; (2) identified pathogen-associated changes in microbiome structure and diversity; (3) elucidated bacterial taxa that correlate with *E. ruminantium* infection status; and (4) explored the functional implications of microbiome alterations for tick physiology and pathogen transmission. Our findings advance understanding of the ecological interactions between *E. ruminantium* and *Am. gemma* ticks, with potential implications for developing microbiome-based control strategies for heartwater disease in livestock production.

## MATERIALS AND METHODS

### Study design

The aim of the study was to describe microbiota composition of *E. ruminantium*-infected and pathogen-free *Am. gemma* tissues at the genus level using tick samples from a previous study (15, 26). In April 2024, ticks were collected from cattle in Kajiado County, southern Kenya (see Ethical approval and permits below), and transported live to International Centre of Insect Physiology and Ecology (*icipe*) Martin Lüscher Emerging Infectious Diseases (ML-EID) Laboratory in Nairobi for identification, tissue collection and further molecular analysis. In the laboratory, the ticks were morphologically identified using taxonomic keys (38, 39, 40) and dissected for haemolymph, midgut and salivary gland collection as described by Getange et al. (26) and Khogali et al. (16, 17, 41, 42). High throughput 16S rRNA gene sequencing was used to compare microbial community composition between *E. ruminantium* infected and uninfected tick samples. Alpha- and beta-diversity metrics, differential abundance analysis, and co-occurrence network analysis were performed to evaluate community structure and interactions, including identification of keystone taxa. Functional profiles were predicted using PICRUSt2 to assess potential associations with infection status.

### Extraction of DNA and confirmation of *E. ruminantium* infection status

Genomic DNA was extracted from dissected tick tissues (haemolymph, midgut, and salivary glands) of 23 *Am. gemma* ticks collected from Kajiado County, Kenya. Extractions were performed using the Bioline ISOLATE II Genomic DNA Kit (Meridian Bioscience, Cincinnati, OH, USA) following the manufacturer’s protocol with minor modifications. Briefly, tissues were lysed in proteinase K buffer at 56°C for 4 hours, followed by column-based purification. All procedures were conducted under sterile conditions to minimize contamination risk. Three extraction negative controls consisting of nuclease free water were included in the extraction and processed alongside the tissue samples. DNA concentration and purity were assessed using a NanoDrop spectrophotometer (Thermo Fisher Scientific, Waltham, MA, USA). Infection status was confirmed by conventional PCR using genus-specific primers PER1 (5’-TTTATCGCTATTAGATGAGCCTATG-3’) and PER2 (5’-CTCTACACTAGGAATTCCGCTAT-3’) (43), classifying samples as *E. ruminantium* infected (n=11) or *E. ruminantium* uninfected (n=12). A tick was classified as infected if *E. ruminantium* DNA was detected in at least one tissue (haemolymph, midgut, or salivary glands) by PCR, and uninfected only if all three tested negative.

### Library preparation and Nanopore sequencing

A near-full-length bacterial 16S rRNA gene (∼1,500 bp) was amplified using universal primers 27F (5’-AGAGTTTGATCCTGGCTCAG-3’) (44) and 1492R (5’-GGTTACCTTGTTACGACTT-3’) (45). Each 20-µL PCR reaction contained 10 µL of LongAmp Taq 2X Master Mix (NEB, M0287; New England Biolabs, Ipswich, MA, USA), 1 µL each of forward and reverse primer (10 µM), 10 ng of template DNA quantified with the Qubit 1X dsDNA HS Assay Kit (Thermo Fisher Scientific) on a Qubit 4 Fluorometer, and the volume brought up to 20 µL using nuclease-free water. Cycling conditions were as follows: initial denaturation at 95°C for 1 min; 40 cycles of 95°C for 20 s, 55°C for 30 s, and 65°C for 2 min; and a final extension at 65°C for 5 min, followed by a hold at 4°C. The cycle number was set to 40 to improve amplification yield from low-biomass tick tissue DNA.

Sequencing libraries were prepared using the Oxford Nanopore Technologies (ONT) Native Barcoding Kit 96 V14 (SQK-NBD114.96; ONT, Oxford, UK) according to the manufacturer’s protocol. Amplicons were quantified using Qubit, pooled in equimolar ratios, and purified with AMPure XP Beads (Beckman Coulter, Indianapolis, IN, USA) at a 1:1 ratio. Barcoded libraries were then pooled at 50 fmol prior to adapter ligation. The final pooled libraries, comprising 69 biological samples, three extraction negative controls, and two library-preparation negative controls containing nuclease-free water, were loaded onto an R9.4.1 flow cell (FLO-MIN106D; ONT) and sequenced for 72 h on a MinION Mk1B (ONT) using MinKNOW for data acquisition.

### Bioinformatics processing of Nanopore 16S rRNA sequences

Raw FAST5 files from MinKNOW were basecalled with Guppy v6.5.7 (ONT) using the super-accurate model (dna_r9.4.1_450bps_sup.cfg) and demultiplexed with Guppy Barcoder using the Native Barcoding Kit 96 V14, requiring valid barcodes at both read ends. Demultiplexed FASTQ files were imported into QIIME 2 v2023.5 (46) for downstream processing. Read quality was assessed using NanoPlot v1.41 (47). NanoFilt v2.8 (47) was used for quality filtering, retaining reads of 800–1,500 bp with mean quality ≥ Q10. Primers were removed with Cutadapt v3.5 (48; error tolerance 0.1, minimum overlap 10 bp). Reads were dereplicated and screened for chimeras using VSEARCH v2.22.1 (49) against the SILVA 138.1 database (50). Non-chimeric sequences were clustered into operational taxonomic units (OTUs) at 90% sequence similarity using VSEARCH, a threshold appropriate for genus-level resolution given R9.4.1 read accuracy (85–94%) (51). Taxonomic classification was assigned using a naïve Bayes classifier trained on the SILVA 138 99% OTU reference database (50), implemented via the q2-feature-classifier plugin (52). Resulting feature, taxonomy, and sequence tables from the QIIME2 pipeline were exported for statistical analysis in R v4.3.1 (53). Rarefaction curves were generated to assess sequencing depth. OTUs classified as mitochondria, chloroplasts, Archaea, Cyanobacteria, Chloroflexi, Eukaryota, or unassigned at the kingdom level were excluded. Potential contaminant sequences were identified by comparing prevalence in true samples versus negative controls using the *decontam* package (54) in R (53), with exclusion of taxa scoring ≥ 0.5.

### Tissue microbiome analysis

Alpha diversity metrics were estimated from unrarefied genus-level count data using the phyloseq package (55) in R. Three alpha diversity indices were calculated: observed richness, Shannon diversity index (56), and Pielou’s evenness (57). Differences in alpha diversity among tick tissue infection status groups were assessed using Kruskal–Wallis tests (α = 0.05), with post hoc pairwise comparisons performed using Wilcoxon rank-sum tests with Bonferroni correction. Beta diversity was assessed using the Bray–Curtis dissimilarity index (58), calculated from relative abundance-transformed genus-level count data, which accounts for differences in sequencing depth across samples. Community composition was visualised using Principal Coordinates Analysis (PCoA). Differences in community composition between groups were tested using permutational multivariate analysis of variance (PERMANOVA; 9,999 permutations, *p* < 0.05), implemented via the adonis2 function in the vegan package (59). To verify that PERMANOVA results reflected true compositional differences rather than differences in within-group dispersion, homogeneity of multivariate dispersions was assessed using the betadisper function in vegan, with significance evaluated by permutation test. Relative abundances were visualised using stacked bar plots of the 15 most abundant genera, pooled by tissue–infection status group and constructed in ggplot2 (60). Genera outside the top 15 were collapsed into an “others” category. Mean relative abundances per genus within each group were additionally used to construct heatmaps using the ComplexHeatmap package (61), with columns stratified by *Ehrlichia* infection status to facilitate direct comparison across tissue types.

### Bacterial co-occurrence networks and statistical network estimation

Co-occurrence networks were constructed separately for each tissue–infection status group using genus-level taxonomic profiles. Pairwise correlations were estimated using the SparCC method (62) implemented through the NetCoMi package (63) in R. Both positive (≥ 0.5) and negative (≤ −0.5) correlations were retained, and the resulting adjacency matrices were exported for visualization and further analysis. Zeros were handled using multiplicative replacement prior to correlation estimation. Network visualization and topological characterisation, including number of nodes and edges, network diameter, modularity, average degree, weighted degree, clustering coefficient, and number of modules were performed in Gephi v0.10 (64).

### Differential network analysis

To compare networks between *E. ruminantium*-infected and *E. ruminantium*-free samples within each tissue type, a structural similarity analysis was conducted using the NetCoMi package (63) implemented in R. To test for dissimilarities between paired networks, the Jaccard index was calculated for unweighted degree, weighted degree, betweenness centrality, and eigenvector centrality. The Jaccard index quantifies overlap between sets of central nodes (centrality above the 75th percentile) and ranges from 0 (no overlap) to 1 (identical), with significance assessed via P(J ≤ j) and P(J ≥ j) computed against the total taxa shared between networks (63). Permutation tests with 1,000 iterations were used to determine statistical significance. Hub taxa similarity was additionally assessed by comparing the top 10 nodes ranked by eigenvector centrality between each paired group.

### Keystone taxa identification

Keystone taxa were defined using three criteria as previously reported (26, 65): (i) ubiquity – microbial taxa present in 100% of samples within a given tissue–infection status group; (ii) eigenvector centrality greater than 0.75; and (iii) mean CLR abundance exceeding the mean abundance of all taxa within the same group. Eigenvector centrality values were exported from Gephi v0.10 (64). Genus-level count data were transformed using centred log-ratio (CLR) normalisation via the microbiome package (66) to account for compositional constraints, and mean CLR values were calculated for each taxon within each group. Taxa meeting only two of the three criteria were additionally identified as candidate keystones. Mean CLR abundance and eigenvector centrality values were plotted as scatter plots using ggplot2 (59) in R (53).

### Functional prediction and differential analysis

The functional potential of microbial communities was predicted from 16S rRNA gene sequences using PICRUSt2 (67), executed via the standalone picrust2_pipeline.py script with representative sequences (seqs.fna) and the OTU table (table.biom) as inputs. Functional annotations were added using add_descriptions.py. OTUs were placed into a reference phylogenetic tree containing more than 20,000 full-length prokaryotic 16S rRNA sequences, applying a nearest-sequenced taxon index (NSTI) cut-off of 3. The use of near-full-length 16S rRNA gene sequences from Nanopore sequencing provides higher-resolution phylogenetic placement than short-read amplicons (e.g., V1–V2 or V3–V4 regions), improving the accuracy of PICRUSt2’s reference-tree-based functional inferences. Quality control analysis confirmed that all samples passed the NSTI threshold with weighted values < 0.15 (mean ± SD: 0.09 ± 0.02). Gene family copy numbers were predicted per OTU based on KEGG Ortholog (KO) annotations (68) and Enzyme Commission (EC) numbers, with metabolic pathway reconstruction performed against the MetaCyc database (69).

Differential abundance of predicted functional pathways was assessed in R (53) by comparing *Ehrlichia*-positive and -negative samples within each tissue type using DESeq2 (70). Significant functional differences were identified at a false discovery rate (FDR) threshold of *p* < 0.05. Results were visualised as scatter plots, volcano plots, and heatmaps of the top differentially abundant pathways and enzyme functions, generated using ggplot2 (59) and pheatmap (71).

## RESULTS

### Taxonomic composition and diversity of *Am. gemma* tissue microbiota

A total of 74 tick samples (23 haemolymph, 23 midguts, 23 salivary glands and 5 negative controls) were sequenced using Oxford Nanopore 16S rRNA gene amplicon sequencing. From an initial total of 1.4 million raw reads obtained via Oxford Nanopore sequencing of a near-full 16S rRNA gene, a robust dataset of 672,322 reads remained after stringent quality filtering and the removal of non-target DNA. After further quality filtering to remove unassigned taxa, six tissue samples from two ticks were excluded due to insufficient sequencing depth (<1,000 reads after decontamination), retaining a total of 339,266 reads from 63 samples across 21 ticks for downstream analysis (**Table S1**). These yielded 1,084 operational taxonomic units (OTUs), which aggregated into 15 phyla, 23 classes, 70 orders, 119 families, and 226 genera. All five negative controls yielded less than 50 reads after filtering suggesting minimal contamination.

Tick microbiome was dominated by Proteobacteria, followed by Firmicutes, Bacteroidota, and Actinobacteriota at the phylum level (Figure 1). The most abundant genera were *Coxiella*, *Pseudomonas*, *Acinetobacter*, *Proteus*, and *Rickettsia* (**Table 1**; **Figure 2**). *Ehrlichia ruminantium* was recovered at 2.51% mean relative abundance. Other prevalent genera included *Staphylococcus*, *Paracoccus*, *Sphingobacterium*, *Gemmobacter*, *Streptococcus*, *Paenibacillus*, and *Serratia*, highlighting a complex microbial community consisting of both environmental and host-associated bacteria (**Table 1**; **Figure 1A**). Notably, multiple tick-associated endosymbionts and pathogens were identified, including *Coxiella*, *Rickettsia*, *Ehrlichia*, *Francisella*, *Wolbachia*, and *Candidatus* Midichloria, consistent with known tick symbiotic and pathogenic bacterial taxa. Notably, *Rickettsia* was consistently more abundant across all three tissues in uninfected ticks, whereas *Ehrlichia* was detected predominantly in the midgut and salivary glands of infected ticks, suggesting tissue-specific colonisation dynamics. Rarefaction curves approached saturation for most samples, with gradual increases in some indicating residual low-abundance taxa (**Figure S1**).

**Figure 1.**
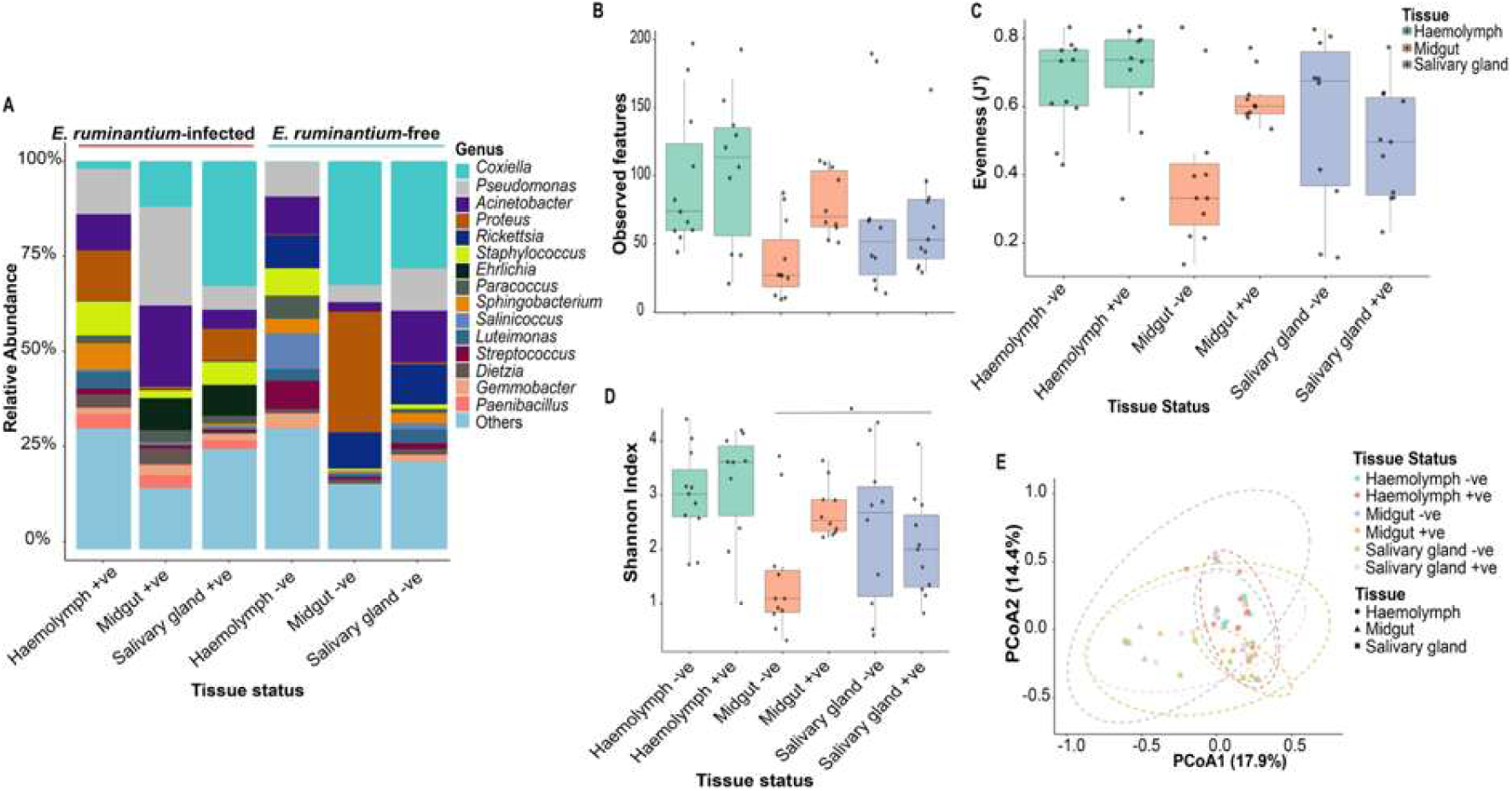
Microbial community composition and diversity of *Ehrlichia ruminantium*-positive and -negative *Amblyomma* tick tissues. **(A)** Relative abundance heatmap of the top 15 bacterial genera pooled by tissue and pathogen status. Alpha diversity by tissue and pathogen status. **(B)** Observed richness. **(C)** Pielou’s evenness index. **(D)** Shannon index. **(E)** Principal coordinates analysis (PCoA) of Bray–Curtis dissimilarity at the genus level showing clustering by tissue and pathogen status.

**Figure 2.**
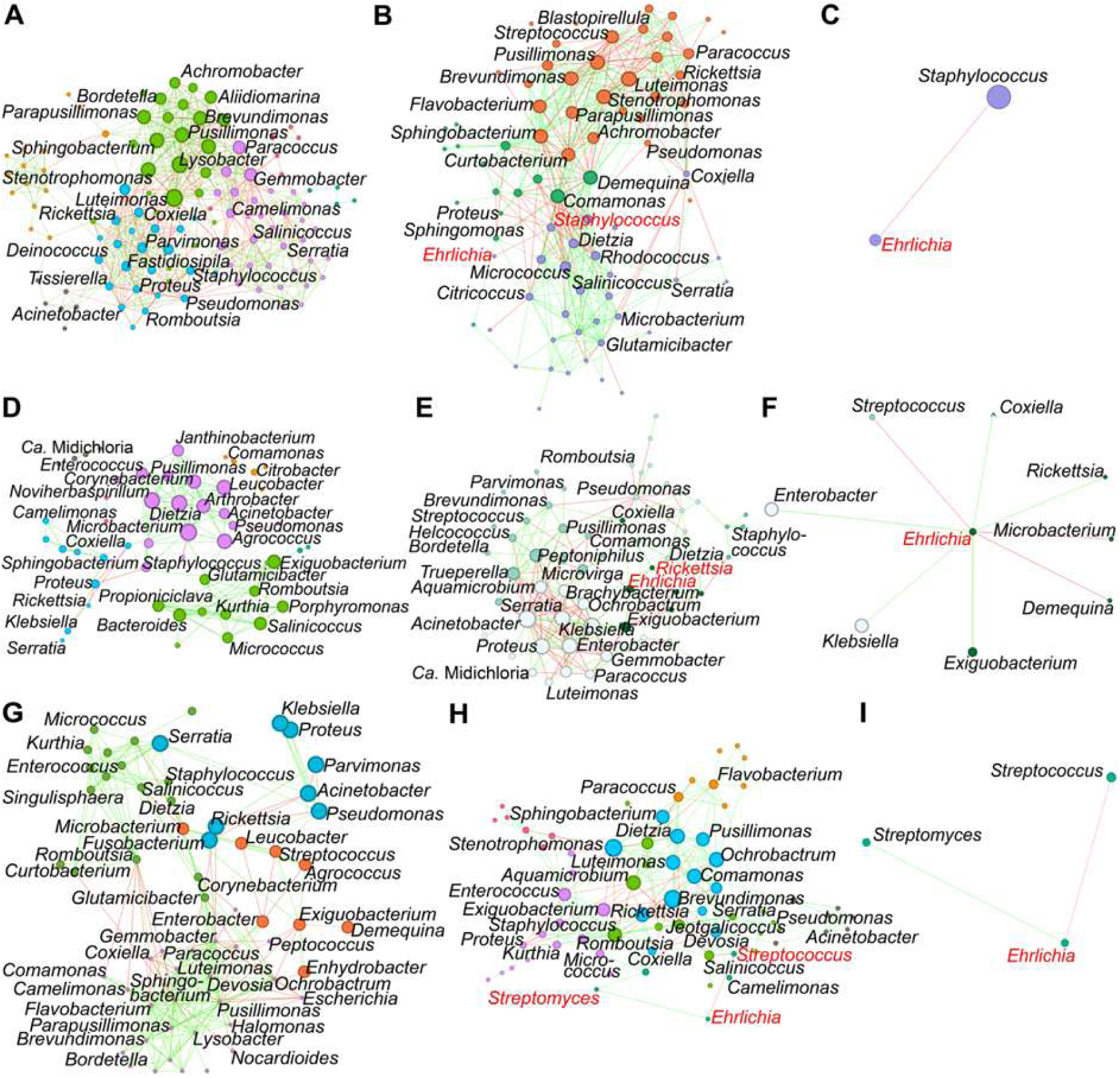
Tissue-specific bacterial co-occurrence networks in *Amblyomma gemma* ticks by *Ehrlichia ruminantium* infection status. Co-occurrence networks of **(A)** *E. ruminantium*-free and **(B)** *E. ruminantium*-infected haemolymph samples. The direct neighbours of *E. ruminantium* were identified in the haemolymph co-occurrence networks and represented as a sub-network **(C)**. Co-occurrence networks of **(D)** pathogen-free and **(E)** *E. ruminantium*-positive midgut samples, with the corresponding *E. ruminantium* sub-network **(F)**. Co-occurrence networks of **(G)** pathogen-free and **(H)** *E. ruminantium*-positive salivary gland samples, with the corresponding *E. ruminantium* sub-network **(I)**. Nodes correspond to bacterial taxa at genus level, and connecting edges indicate significant co-occurrence relationships between them. Only nodes with at least one significant correlation are shown. Node size is proportional to eigenvector centrality, and node colour reflects modularity class, whereby taxa of the same colour belong to the same co-occurrence module. Edge colours represent strong positive (green) or negative (red) correlations (SparCC ≥ 0.5 or ≤ −0.5).

**Table 1.** Mean relative abundance (%) of the top 15 bacterial genera across *Ehrlichia ruminantium*-infected and *E. ruminantium*-free tick tissues.

| Genus | *E. ruminantium* infection |  |  | *E. ruminantium*-free |  |  |
| --- | --- | --- | --- | --- | --- | --- |
|  | Haemolymph (%) | Midgut (%) | Salivary gland (%) | Haemolymph (%) | Midgut (%) | Salivary gland (%) |
| *Coxiella* | 2.053 | 11.694 | 32.196 | 0.104 | 31.942 | 27.608 |
| *Pseudomonas* | 11.569 | 25.462 | 6.063 | 8.997 | 4.366 | 10.906 |
| *Acinetobacter* | 9.374 | 20.922 | 4.871 | 9.682 | 2.432 | 13.33 |
| *Proteus* | 13.014 | 0.708 | 8.271 | 0.249 | 31.154 | 0.459 |
| *Rickettsia* | 0.115 | 0.265 | 0.265 | 8.542 | 9.303 | 10.353 |
| *Staphylococcus* | 8.748 | 1.971 | 5.929 | 7.003 | 0.477 | 1.121 |
| *Ehrlichia* | 0.055 | 8.316 | 7.98 | 0.007 | 0.002 | 0 |
| *Paracoccus* | 1.976 | 3.077 | 2.081 | 6.04 | 0.056 | 1.1 |
| *Sphingobacterium* | 6.754 | 0.095 | 0.592 | 3.69 | 0.408 | 2.644 |
| *Salinicoccus* | 0.548 | 0.482 | 0.511 | 9.189 | 0.474 | 1.55 |
| *Luteimonas* | 4.409 | 0.249 | 0.489 | 3.088 | 0.707 | 3.55 |
| *Streptococcus* | 1.404 | 0.735 | 0.627 | 7.261 | 0.601 | 1.542 |
| *Dietzia* | 3.509 | 4.233 | 0.333 | 1.117 | 1.179 | 1.477 |
| *Gemmobacter* | 1.556 | 2.705 | 1.689 | 3.806 | 0.019 | 1.702 |
| *Paenibacillus* | 3.772 | 3.296 | 2.294 | 0.122 | 0.022 | 0.044 |

All three alpha-diversity indices (Shannon, observed richness, Pielou’s evenness) differed significantly among tissue-status groups (Kruskal–Wallis *p* ≤ 0.011) (**Figure 1B-D**; **Table 2**). Pairwise comparisons revealed that midgut negative samples had significantly lower Shannon diversity compared to haemolymph negative samples (*p* = 0.037), with no other significant pairwise differences after Bonferroni correction (**Table 3**). Uninfected midguts showed the lowest diversity across all metrics, while haemolymph samples showed the highest, regardless of *Ehrlichia* infection status (**Table 4**). Bray–Curtis PCoA revealed distinct clustering by tissue–infection group, with PERMANOVA confirming a strong group effect (F = 1.89, R² = 0.142, *p* < 0.001; Figure 1E) and homogeneous dispersion (betadisper *p* = 0.134). Pairwise FDR-corrected PERMANOVA identified differences between uninfected haemolymph vs. midgut (*p* = 0.015) and uninfected vs. infected midgut (*p* = 0.030; **Table 3**).

**Table 2.** Overall microbial diversity statistics.

| Diversity Metric | Test | $\chi^2$ Statistic | df | p-value |
| --- | --- | --- | --- | --- |
| Shannon Diversity | Kruskal-Wallis | 16.284 | 5 | **0.006** |
| Observed Richness | Kruskal-Wallis | 15.077 | 5 | **0.010** |
| Pielou's Evenness | Kruskal-Wallis | 14.868 | 5 | 0.011 |
| Beta Diversity | PERMANOVA | F = 1.886 | 5, 57 | **< 0.001** |

**Table 3.** Significant pairwise comparisons between tissue-status groups.

| Analysis | Comparison | Test Statistic | Raw p-value | Adjusted p-value |
| --- | --- | --- | --- | --- |
| Shannon Diversity | MDN vs HLN | — | 0.037 | **0.037\*** |
| Beta Diversity | HLN vs MDN | F = 3.05 | 0.001 | **0.015\*** |
| Beta Diversity | MDN vs MDP | F = 4.22 | 0.002 | **0.030\*** |

HLN: Haemolymph pathogen-free; MDN: Midgut pathogen free; MDP: Midgut *Ehrlichia* infection

**Table 4.** Alpha diversity summary by tissue-status group.

| Group | *Ehrlichia* Status | n | Shannon (mean ± SD) | Observed (mean ± SD) | Pielou's Evenness (mean ± SD) |
| --- | --- | --- | --- | --- | --- |
| Haemolymph | Negative | 11 | 3.01 ± 0.85 | 96.6 ± 52.5 | 0.666 ± 0.134 |
|  | Positive | 10 | 3.18 ± 1.06 | 104.5 ± 55.0 | 0.692 ± 0.158 |
| Midgut | Negative | 11 | 1.45 ± 1.11 | 37.5 ± 28.5 | 0.398 ± 0.220 |
|  | Positive | 10 | 2.71 ± 0.50 | 79.3 ± 23.9 | 0.623 ± 0.074 |
| Salivary gland | Negative | 11 | 2.35 ± 1.43 | 70.6 ± 64.4 | 0.555 ± 0.259 |
|  | Positive | 10 | 2.04 ± 0.93 | 65.9 ± 39.3 | 0.488 ± 0.167 |

### Impact of *E. ruminantium* infection on microbial community structure in *Am. Gemma*

To investigate whether *E. ruminantium* infection alters tick microbiota structure, bacterial co-occurrence patterns were quantified using network topology analysis across haemolymph, midgut, and salivary gland tissues. Visually, *E. ruminantium*-free networks across all tissues featured well-connected modules with strong positive interactions between taxa, whereas infected networks displayed notable restructuring in module organisation and interaction patterns (**Figure 2A–F**). Haemolymph networks were the most complex; infection reduced node count (113 vs. 78) and edge count (616 vs. 422) but increased density (0.097 vs. 0.141), suggesting more constrained interactions among fewer taxa (**Table 5**). In midgut networks, hub taxa shifted from environmental genera (*Agrococcus*, *Noviherbaspirillum*, *Dietzia*) to opportunistic Proteobacteria (*Acinetobacter*, *Proteus*, *Klebsiella*, *Serratia*) on infection (**Table S2**), alongside an order-of-magnitude increase in negative edges (7 vs. 77). Salivary gland networks showed intermediate restructuring, with partially conserved hub taxa but altered edge distributions and modularity (**Figure 2A–F**; **Table 5**).

**Table 5.** Co-occurrence network topological properties of bacterial communities across tick tissues and *Ehrlichia ruminantium* infection status.

| Network Property | HLN | HLP | MDN | MDP | SGN | SGP |
| --- | --- | --- | --- | --- | --- | --- |
| **Nodes** | 113 | 78 | 53 | 64 | 60 | 69 |
| **Edges** | 616 | 422 | 131 | 187 | 258 | 223 |
| **Positive edges** | 457 | 309 | 124 | 110 | 208 | 169 |
| **Negative edges** | 159 | 113 | 7 | 77 | 50 | 54 |
| **Network density** | 0.097 | 0.141 | 0.095 | 0.093 | 0.146 | 0.095 |
| **Modularity** | 0.526 | 0.388 | 0.588 | 0.430 | 0.442 | 0.429 |
| **Network diameter** | 8 | 5 | 9 | 7 | 5 | 8 |
| **Average degree** | 10.903 | 10.821 | 4.943 | 5.937 | 8.600 | 6.464 |
| **Av. weighted degree** | 6.47 | 6.176 | 2.840 | 3.525 | 5.106 | 3.794 |
| **Clustering co-efficient** | 0.567 | 0.606 | 0.553 | 0.516 | 0.673 | 0.537 |

HLN, haemolymph negative; HLP, haemolymph positive; MDN, midgut negative; MDP, midgut positive; SGN, salivary gland negative; SGP, salivary gland positive.

Jaccard similarity indices confirmed tissue-specific network reorganisation associated with *E. ruminantium* infection. While degree-based centrality measures were conserved across all tissues (Jaccard = 1.0), hub taxa composition revealed the most pronounced infection-driven changes, with complete replacement of hub taxa in the midgut (Jaccard = 0.000, *p* = 0.043) and partial conservation in haemolymph (Jaccard = 0.429) and salivary glands (Jaccard = 0.177). Modular reorganisation assessed by the Adjusted Rand Index was most significant in salivary glands (ARI = 0.194, *p* < 0.001), while haemolymph and midgut showed non-significant modular shifts (**Tables 6–9**). Sub-network analysis of *E. ruminantium* connectivity revealed tissue-specific interaction patterns: in haemolymph, the pathogen associated negatively with *Staphylococcus* only; in the midgut, it showed broader connectivity with negative associations with *Demequina*, *Microbacterium*, and *Streptococcus*, and positive co-occurrences with *Coxiella*, *Enterobacter*, *Klebsiella*, *Exiguobacterium*, and *Rickettsia*; and in salivary glands, it associated positively with *Streptomyces* and negatively with *Streptococcus*. The midgut positive association between *E. ruminantium* and *Rickettsia*, both obligate intracellular bacteria, should be interpreted cautiously, since networks reflect correlated variation rather than absolute abundance. Although low-level co-detection drives the correlation, overall *Rickettsia* abundance was near-absent in infected tissues (0.3% vs. 9.3% in midgut), consistent with competitive exclusion at the community level. These findings indicate that *E. ruminantium* infection is associated with significant tissue-specific restructuring of microbial community structure, with the midgut emerging as the primary site of network reorganisation.

**Table 6.**
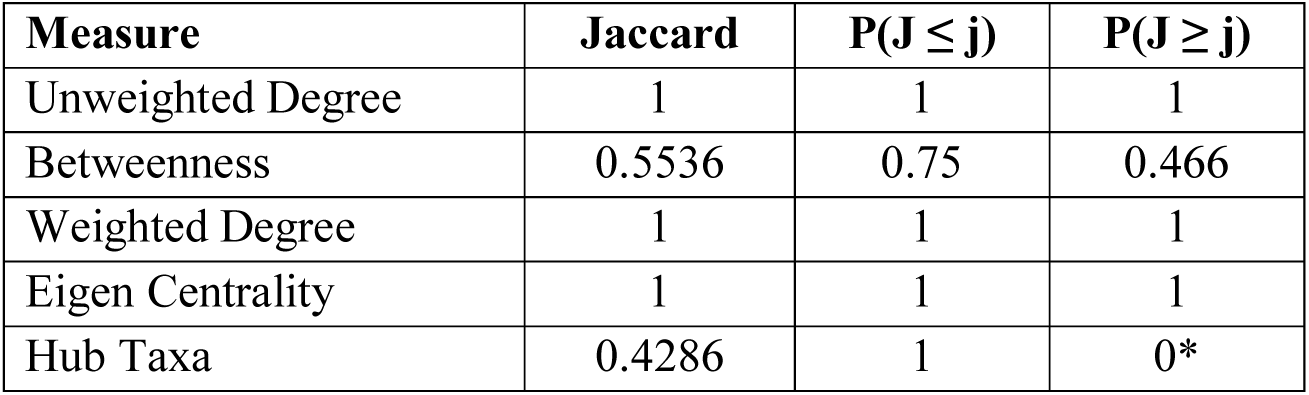
Jaccard index for haemolymph pathogen-free and *E. ruminantium* infection networks.

**Table 7.**
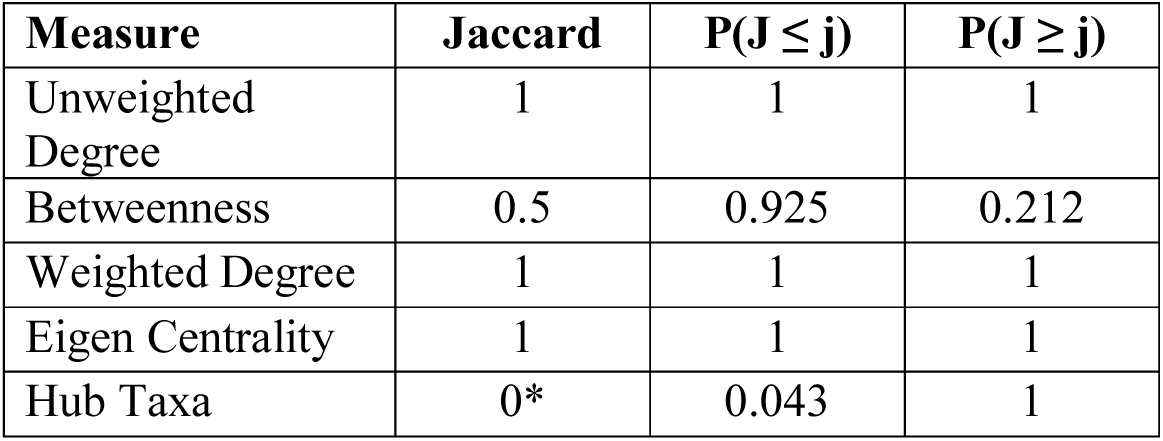
Jaccard index for midgut pathogen-free and *E. ruminantium* infection networks.

| Measure | Jaccard | P(J ≤ j) | P(J ≥ j) |
| --- | --- | --- | --- |
| Unweighted Degree | 1 | 1 | 1 |
| Betweenness | 0.5 | 0.925 | 0.212 |
| Weighted Degree | 1 | 1 | 1 |
| Eigen Centrality | 1 | 1 | 1 |
| Hub Taxa | 0* | 0.043 | 1 |

**Table 8.** Jaccard index for salivary gland pathogen-free and *E. ruminantium* infection networks.

| Measure | Jaccard | P(J ≤ j) | P(J ≥ j) |
| --- | --- | --- | --- |
| Unweighted Degree | 1 | 1 | 1 |
| Betweenness | 0.7188 | 0.998 | 0.006* |
| Weighted Degree | 1 | 1 | 1 |
| Eigen Centrality | 1 | 1 | 1 |
| Hub Taxa | 0.1765 | 0.802 | 0.494 |

**Table 9.** Adjusted Rand Index (ARI) comparison of bacterial community clustering between *E. ruminantium*-free and *E. ruminantium*-infected tick tissues. *P*-values were obtained from permutation testing.

| Tissue | Comparison | ARI Value | *p*-value |
| --- | --- | --- | --- |
| Haemolymph | Pathogen-free vs positive | 0.0167 | 0.461 |
| Midgut | Pathogen-free vs positive | 0.0656 | 0.085 |
| Salivary gland | Pathogen-free vs positive | 0.1937 | < 0.001 |

### Impact of *E. ruminantium* infection on microbial hierarchy of *Am. gemma*

To determine whether *E. ruminantium* infection alters the hierarchical organisation of the *Am. gemma* microbiota, keystone taxa were identified based on three criteria: ubiquity across all samples within a group, high eigenvector centrality (> 0.75), and mean CLR abundance above the community mean. In all three pathogen-free tissues, no taxa met the full criteria for keystone status, although several candidate hubs were identified, notably *Luteimonas* in haemolymph, *Agrococcus* and *Noviherbaspirillum* in midgut, and *Sphingobacterium* and *Flavobacterium* in salivary glands. The absence of keystones in pathogen-free samples suggests that healthy tick microbiomes are characterised by inter-individual variability and functional redundancy rather than fixed hierarchical dominance (**Table 10**; **Figure 3**).

**Figure 3.**
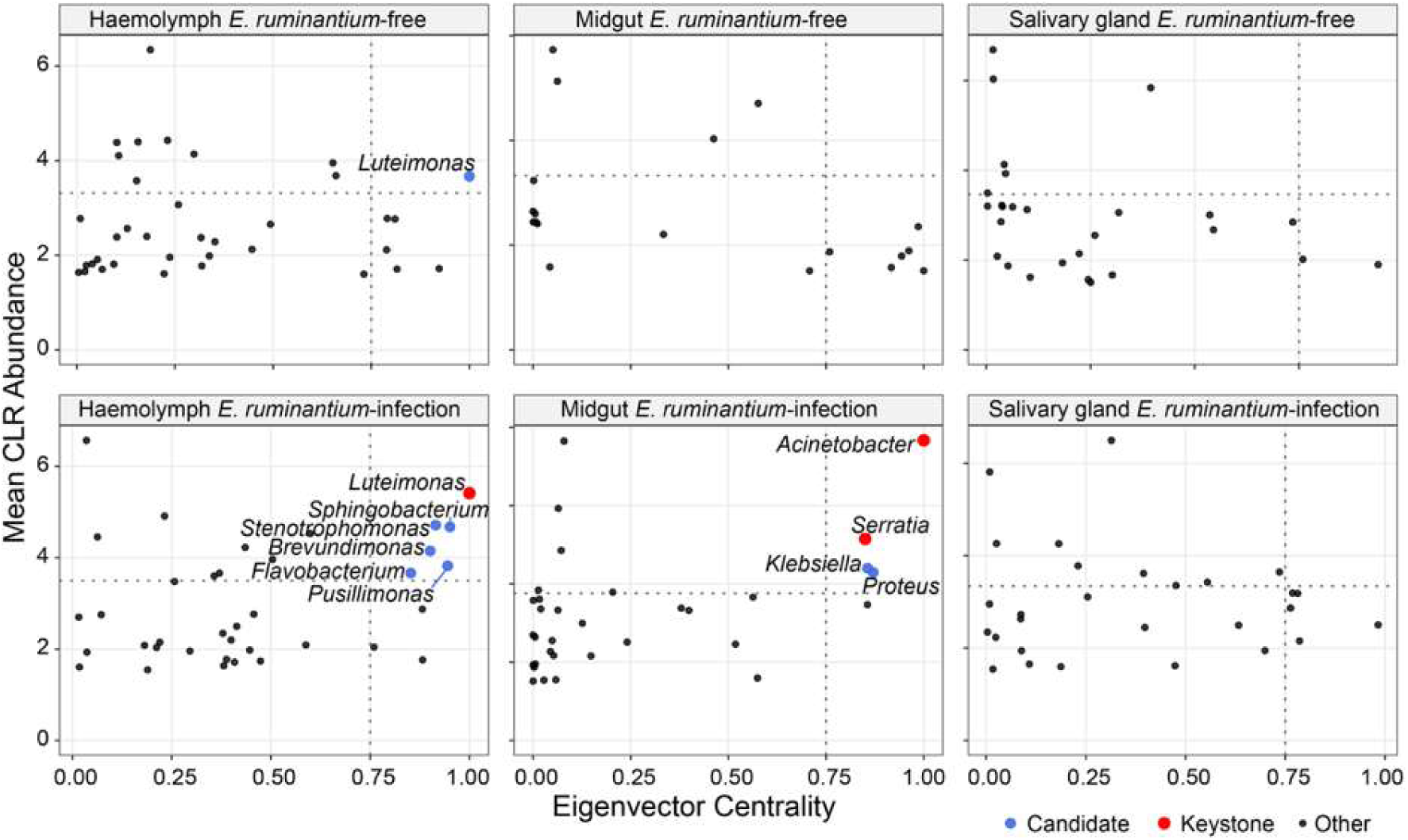
Structural reorganisation of the *Amblyomma gemma* microbiota during *Ehrlichia* infection. Distribution of bacterial genera based on Eigenvector Centrality and Mean CLR abundance (taxa < 2 CLR excluded). Red points denote ubiquitous keystone taxa; blue points indicate non-ubiquitous candidate hubs (Centrality > 0.75; CLR > group mean).

**Table 10.** Comparative network hierarchy and keystone analysis.

| **Tissue** | **Infection status** | **Network state** | **Identified keystone taxa** | **Primary hubs (non-keystones)** |
| --- | --- | --- | --- | --- |
| **Haemolymph** | Negative | Decentralized | None | *Luteimonas*, *Lysobacter* |
|  | Positive | Centralized | *Luteimonas* | (Keystone Dominant) |
| **Midgut** | Negative | Decentralized | None | *Agrococcus*,<br>*Noviherbaspirillum* |
|  | Positive | Centralized | *Acinetobacter*,<br>*Serratia* | (Keystone Dominant) |
| **Salivary glands** | Negative | Decentralized | None | *Sphingobacterium*,<br>*Flavobacterium* |
|  | Positive | Decentralized | None | *Brevundimonas*,<br>*Stenotrophomonas* |

*Ehrlichia ruminantium* infection induced deterministic centralisation of microbial hierarchies in a tissue-specific manner. In infected haemolymph, *Luteimonas* was elevated from candidate hub to keystone status by our prespecified criteria, achieving 100% prevalence and maximum eigenvector centrality (1.0). Infected midgut networks showed emergence of two keystones, *Acinetobacter* and *Serratia*, alongside candidate hubs *Proteus* (centrality = 0.87) and *Klebsiella* (centrality = 0.86), all belonging to Proteobacteria and several recognised as opportunistic pathogens. Salivary gland networks remained decentralised despite infection, with candidate hubs *Brevundimonas* and *Stenotrophomonas* failing to achieve keystone status, potentially reflecting ongoing ecological instability at the primary site of pathogen transmission. The convergence of keystone and candidate hub taxa towards Proteobacteria in infected tissues, in contrast to the phylogenetically diverse hubs observed in pathogen-free samples, suggests that *E. ruminantium* infection is associated with shifts in microbial community hierarchy that may facilitate secondary bacterial colonisation through disruption of competitive exclusion mechanisms (**Table 11**; **Figure 3**).

**Table 11.** Identified candidate hub taxa.

| **Group** | **Status** | **Genus** | **Phylum** | **Mean CLR** | **EigenCentrality** |
| --- | --- | --- | --- | --- | --- |
| **Haemolymph** | Negative | *Luteimonas* | Proteobacteria | 3.67 | 1.000 |
|  | Positive | *Stenotrophomonas* | Proteobacteria | 4.68 | 0.951 |
|  | Positive | *Pusillimonas* | Proteobacteria | 3.82 | 0.945 |
|  | Positive | *Sphingobacterium* | Bacteroidota | 4.72 | 0.915 |
|  | Positive | *Brevundimonas* | Proteobacteria | 4.15 | 0.901 |
|  | Positive | *Flavobacterium* | Bacteroidota | 3.67 | 0.852 |
| **Midgut** | Positive | *Proteus* | Proteobacteria | 4.30 | 0.871 |
|  | Positive | *Klebsiella* | Proteobacteria | 4.40 | 0.856 |

### Impact of *E. ruminantium* infection on the inferred functional profiles of *Am. Gemma* microbiota

PICRUSt2-inferred functional profiles showed substantial core feature sharing between *E. ruminantium*-positive and -negative *Am. gemma* microbiota (85–89% of core pathways, enzymes, and KO functions shared; **Figures 4–6**; **Table 12**). Despite this overlap, Bray–Curtis PCoA revealed significant beta-diversity differences across all three functional layers (PERMANOVA *p* ≤ 0.008; **Figures 4–6**; **Table 13**), with uninfected samples showing higher dispersion (all *p* < 0.001). FDR-significant features confirmed persistent compositional differences for KO functions (*p* = 0.029), supporting genuine biological differences rather than statistical artefacts.

**Figure 4.**
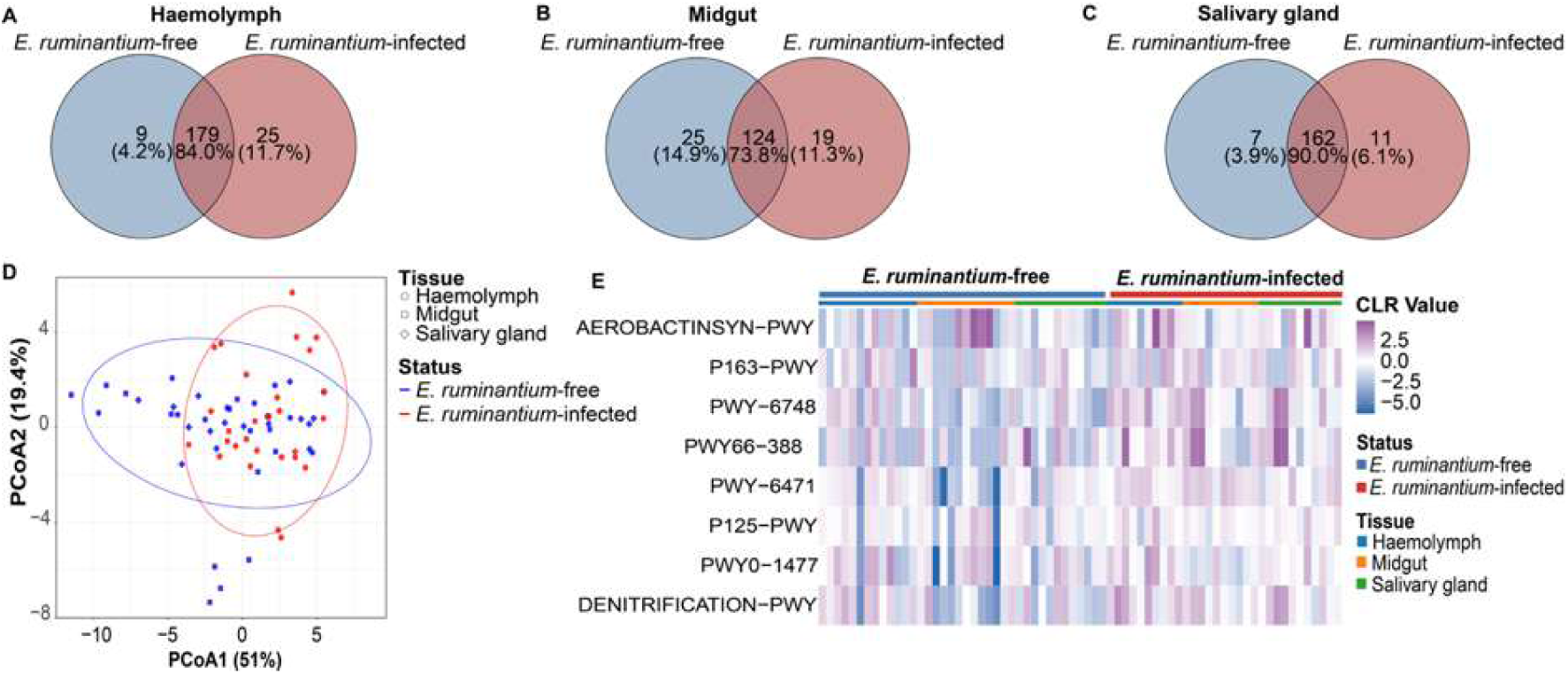
Diversity and differential abundance of predicted metabolic pathways in *Amblyomma gemma* microbiota from *Ehrlichia ruminantium*-infected and *E. ruminantium*-free tissues. Venn diagrams representing unique and shared core metabolic pathways (prevalence ≥75%) of *E. ruminantium*-free (blue) and *E. ruminantium*-infected (red) microbiota in **A.** haemolymph, **B.** midgut, and **C.** salivary gland tissues. **D.** Beta-diversity comparison using Bray–Curtis dissimilarity (PERMANOVA: F₂,₆₆ = 4.66, R² = 0.065, *p* = 0.005). **E.** Heatmap of differentially abundant pathways. Only pathways with statistically significant differences are shown (DESeq2, *p* < 0.05).

**Figure 5.**
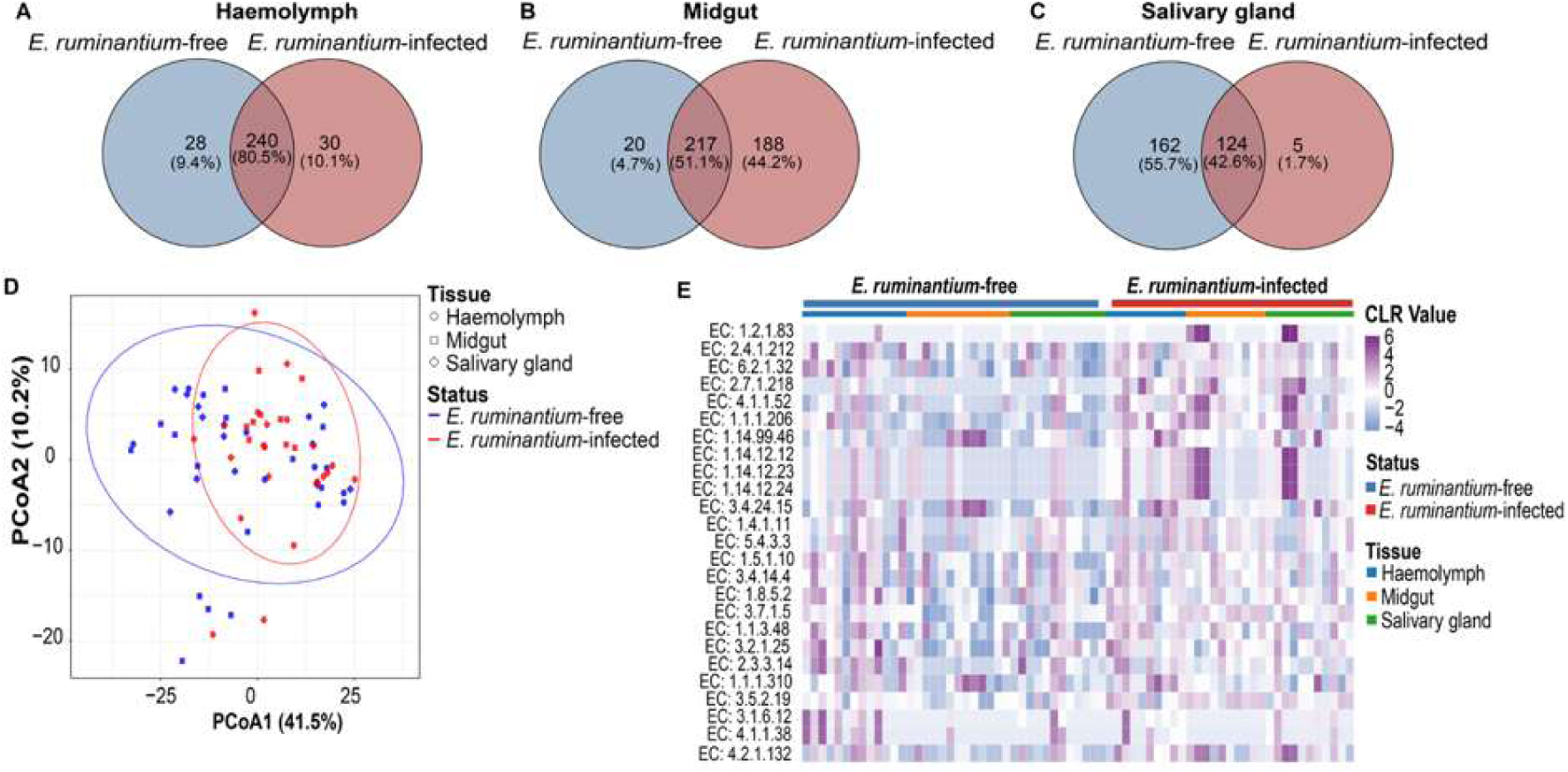
Diversity and differential abundance of predicted enzyme commissions (EC) in *Amblyomma gemma* microbiota from *Ehrlichia ruminantium*-infected and *E. ruminantium*-free tissues. Venn diagrams representing unique and shared core predicted enzymes (prevalence ≥75%) of *E. ruminantium*-free (blue) and *E. ruminantium*-infected (red) microbiota in **A.** haemolymph, **B.** midgut, and **C.** salivary gland tissues. **D.** Beta-diversity comparison using Bray–Curtis dissimilarity (PERMANOVA: F₂,₆₆ = 3.79, R² = 0.054, *p* = 0.008). **E.** Heatmap of differentially abundant enzymes. Only enzymes with statistically significant differences are shown (DESeq2, *p* < 0.05).

**Figure 6.**
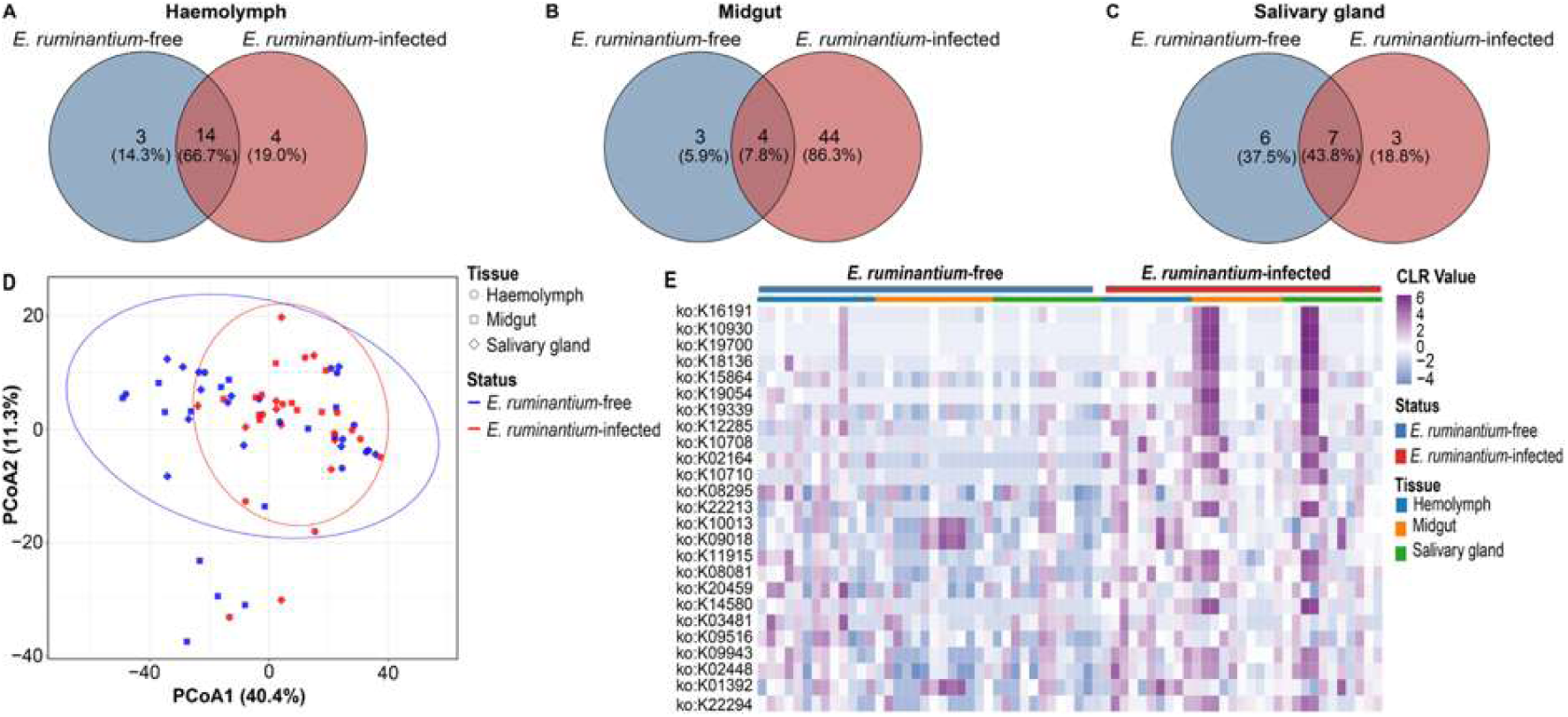
Diversity and differential abundance of predicted gene functions (KEGG Orthology, KO) in *Amblyomma gemma* microbiota from *Ehrlichia ruminantium*-infected and *E. ruminantium*-free tissues. Venn diagrams showing unique and shared core predicted gene functions (prevalence ≥75%) of *E. ruminantium*-free (blue) and *E. ruminantium*-infected (red) microbiota in **A.** haemolymph, **B.** midgut, and **C.** salivary gland tissues. **D.** Beta-diversity comparison using Bray–Curtis dissimilarity (PERMANOVA: F₂,₆₆ = 3.69, R² = 0.052, *p* = 0.007). **E.** Heatmap of differentially abundant gene functions. Only gene functions with statistically significant differences are shown (DESeq2, *p* < 0.05).

**Table 12.**
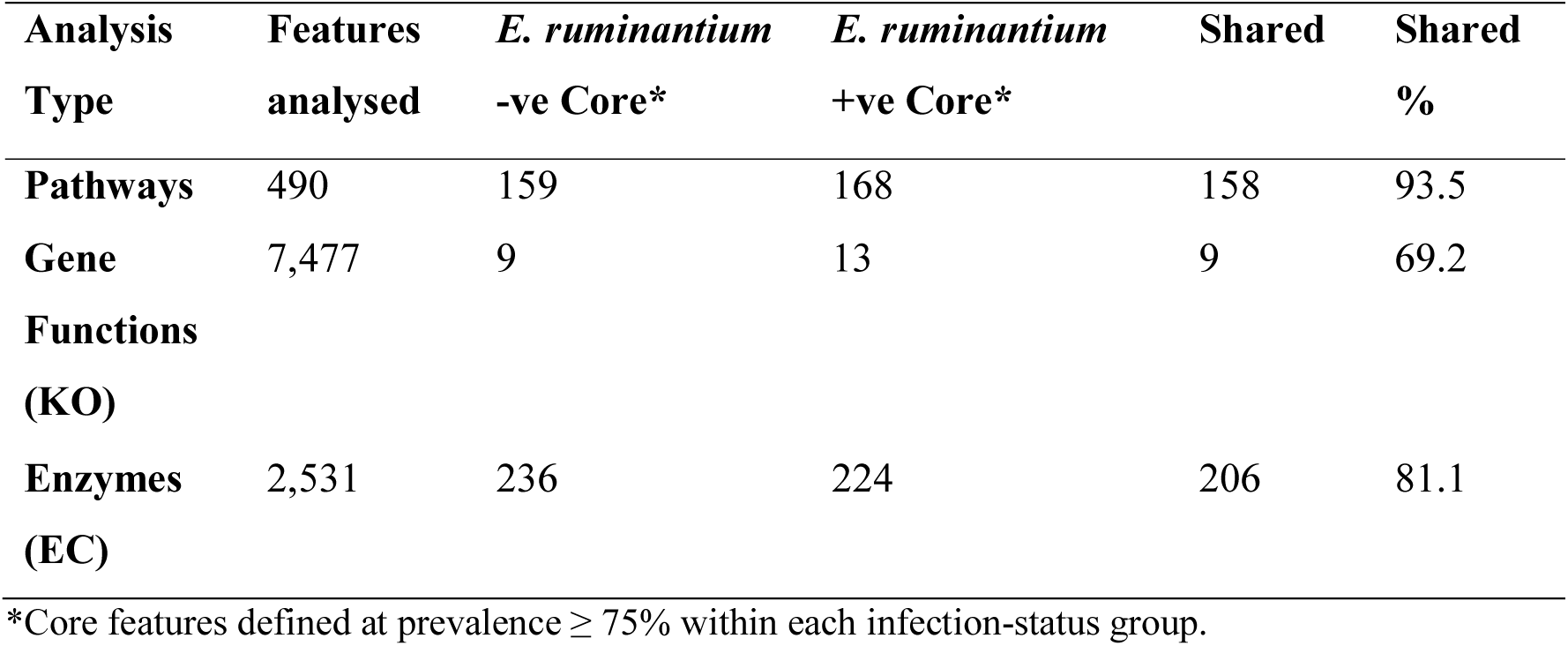
Summary of core functional features shared between *E. ruminantium*-positive and -negative *Am. gemma* microbiota.

**Table 13.** PERMANOVA results for functional beta diversity between *E. ruminantium*-positive and -negative samples.

| Analysis | PERMANOVA F | R<sup>2</sup> | *p* | Dispersion F | Dispersion *p* | *p* < 0.05 | FDR < 0.05 |
| --- | --- | --- | --- | --- | --- | --- | --- |
| **Pathways** | 4.66 | 0.065 | 0.005 | 13.27 | < 0.001 | 8 | 0 |
| **EC enzymes** | 3.79 | 0.054 | 0.008 | 14.41 | < 0.001 | 149 | 2 |
| **KO functions** | 3.69 | 0.052 | 0.007 | 17.14 | < 0.001 | 364 | 9 |

"*p* < 0.05" = unadjusted threshold; "FDR < 0.05" = Benjamini–Hochberg correction.

DESeq2 differential abundance analysis after multiple testing correction (FDR < 0.05) identified 0 pathways, 2 enzymes, and 9 KO functions significantly enriched in *E. ruminantium*-positive samples (**Tables 12, 13**; **Figures 4–6**). The two FDR-significant enzymes were aldehyde dehydrogenase (EC 1.2.1.83; log₂FC = 7.23), involved in oxidative stress detoxification, and LPS glucosyltransferase (EC 2.4.1.212; log₂FC = −2.96), depleted in infected samples and associated with outer membrane remodelling (**Table S3; Figure 5E**). The 9 FDR-significant KO functions converged on two themes: Iron acquisition functions (aerobactin siderophore biosynthesis/transport K16191, K18136, K15864, K19054 plus iron/manganese ABC transport K10708) (log₂FC = 3.02–5.98) and antimicrobial efflux mechanisms (K10930, K19700, K19339, K12285; log₂FC = 2.80–7.32), suggesting robust enrichment of iron acquisition and antimicrobial resistance systems in infected microbiota (**Table S4; Figure 6E**). At the nominal significance threshold (p < 0.05, unadjusted), a broader but exploratory set of 8 pathways, 149 enzymes, and 364 KO functions showed differences (**Table 13**; **Figures 4–6E**), extending the above themes to include amino acid fermentation (P163-PWY; log₂FC = −2.14) and denitrification pathways (log₂FC = 0.83; **Figure 4E**). Among nominally significant pathways, aerobactin biosynthesis (AEROBACTINSYN-PWY) showed a negative pathway-level log₂FC (−2.56) despite positive enrichment of individual aerobactin genes, a discordance that likely reflects taxon-restricted upregulation rather than community-wide enrichment (see Discussion). These nominally significant findings should be regarded as hypothesis-generating pending replication with larger sample sizes.

## DISCUSSION

The capacity of TBPs to modulate the diversity, bacterial composition, and microbial structure of the tick host microbiota has been established across multiple tick species (31, 33–35, 72). However, their impact on microbial community assembly, network properties, and metabolic function, particularly for *Am. gemma* tick, remains poorly understood. In this study, we examined how *E. ruminantium* infection alters the bacterial microbiota of *Am. gemma* tick tissues using Oxford Nanopore near-full-length 16S rRNA gene amplicon sequencing, co-occurrence network analysis, and functional profiling. This combination of tissue-resolved sampling, near-full-length 16S sequencing, and network-based analyses resolves patterns that would be averaged out in whole-tick studies and obscured by short-read taxonomic resolution. Uninfected tissues showed dispersed microbial networks dominated by *Coxiella*, *Pseudomonas*, and endosymbiotic *Rickettsia*, without clear keystone taxa. While baseline tick microbiomes exhibit functional redundancy that may buffer against minor perturbations, our findings suggest that *E. ruminantium* infection may exceed this buffering capacity and produce deterministic restructuring around opportunistic Proteobacteria. Infected tissues converged around specific keystones, notably *Luteimonas* in haemolymph and opportunistic Proteobacteria (*Acinetobacter*, *Serratia*) in midgut, alongside competitive exclusion of *Rickettsia* and functional enrichment of iron acquisition pathways. These tissue-specific variations may suggest that *E. ruminantium* colonisation is associated with disruption of the ecological balance of the *Am. gemma* microbiome, which may reduce colonisation resistance and create conditions permissive to secondary opportunistic colonisation.

*Ehrlichia ruminantium* infection was associated with marked shifts in the *Am. gemma* microbiome, with infected tissues showing higher abundances of *Coxiella*, *Acinetobacter*, *Serratia*, and *Luteimonas*, while *Rickettsia* was near-absent in infected haemolymph, midgut and salivary glands, unlike in uninfected tissues. Similar pathogen-driven abundance shifts have been reported in *Ix. ricinus* and *Ix. scapularis* (33, 35), and Rickettsial taxa have been shown to drive community assembly through privileged connections with keystone taxa despite low network centrality in *Hyalomma marginatum* and *Rhipicephalus bursa* (73). The near-complete absence of *Rickettsia* from the core microbiota of infected samples suggests competitive exclusion consistent with rickettsial exclusion phenomena in *Dermacentor* ticks (36, 37), likely reflecting direct niche competition between obligate intracellular bacteria or indirect competition mediated by altered tick immune responses (36, 74). We note, however, that the cross-sectional design cannot distinguish active exclusion of *Rickettsia* by *E. ruminantium* from preferential establishment of *E. ruminantium* in ticks with already-low *Rickettsia* loads; both mechanisms are consistent with the observed pattern. The positive midgut network association between *E. ruminantium* and *Rickettsia* does not contradict this interpretation; it likely reflects residual signal in a small subset of infected samples against the dominant population-level trend. Notably, we did not discriminate between endosymbiotic and pathogenic *Rickettsia* lineages, so whether these functional groups respond differently to *E. ruminantium* infection remains an open question requiring species-level resolution.

Co-occurrence network analysis revealed that *E. ruminantium* infection may destabilise the microbial community structure in *Am. gemma*, making it more susceptible to disruptions and highlighting tissue-specific responses to pathogen infection. We observed high modularity in uninfected tissue networks and substantially reduced modularity and altered connectivity in *E. ruminantium*-infected networks, suggesting disruption of balanced microbial associations that potentially creates a more pathogen-permissive environment. Similarly to previous findings with *R. helvetica* in *Ix. ricinus* where pathogenic infection reduced microbial connectivity (33), our study indicates that *E. ruminantium* infection alters network structure through tissue-specific mechanisms. Infected haemolymph networks showed increased density alongside fewer connections, while midgut networks showed the greatest reorganisation with increased negative edges, reflecting intense microbial competition. These findings align with previous studies demonstrating compromised network robustness in *A. phagocytophilum*-infected *Ix. scapularis* (35) and microbiota modulation by *A. marginale* in *Rh. microplus* organs involved in pathogen transmission (34). Combined, these findings suggest that various TBPs drive tissue-specific microbial restructuring where network-level changes may facilitate pathogen colonisation or represent downstream consequences of established infection, with the midgut serving as the primary site of microbial community reorganisation. These tissue-specific patterns are consistent with the dual selective pressure framework proposed by Piloto-Sardiñas et al. (75), in which baseline ecological filters and pathogen-driven selection interact to determine microbial network configuration within the tick holobiont.

Our study findings indicate that *E. ruminantium* infection is associated with broad functional conservation in the *Am. gemma* microbiome, consistent with patterns documented across tick species (35, 76). The clearest infection-associated signal, and the only one surviving multiple testing correction, centred on iron acquisition and antimicrobial resistance. Enrichment of aerobactin siderophore biosynthesis genes likely points to intensified competition for iron between the resident microbiota and *E. ruminantium*, an obligate intracellular pathogen that depends on host iron for replication (77), though the cross-sectional sampling does not establish directionality. On the other hand, the concurrent depletion of LPS glucosyltransferase (EC 2.4.1.212) observed in this study may suggest remodelling of outer membrane composition, which echoes the functional disruptions reported in *R. helvetica*-infected *Ix. ricinus* (33). However, the aerobactin biosynthesis pathway (AEROBACTINSYN-PWY) returned a negative log₂FC (−2.56) despite positive enrichment of individual aerobactin genes, a discordance that likely reflects taxon-restricted upregulation, particularly within the opportunistic Proteobacteria (*Klebsiella*, *Serratia*) that dominated infected midgut networks (a recognised limitation of MetaCyc pathway reconstruction in PICRUSt2). Nominally significant findings (unadjusted *p* < 0.05), spanning amino acid fermentation, denitrification, and efflux systems extend these themes but remain exploratory (34, 78). Whether *E. ruminantium* actively drives these changes, as witnessed in *A. phagocytophilum* through induction of the IAFGP protein in *Ix. scapularis* (35, 79), or whether they arise as downstream consequences of established infection is a question that only experimental work can resolve.

The findings from this study have important implications for developing microbiome-based interventions against heartwater transmission. The identification of tissue-specific keystone taxa, *Luteimonas* in infected haemolymph and *Acinetobacter*/*Serratia* in infected midgut, provides candidate targets for anti-microbiota vaccination strategies designed to disrupt microbial conditions required for efficient *E. ruminantium* colonisation (31, 33). Additionally, the competitive exclusion between *E. ruminantium* and endosymbiotic *Rickettsia* may be exploitable through strategies reinforcing *Rickettsia*-mediated colonisation resistance, paralleling rickettsial exclusion in *Dermacentor* ticks (37). The aerobactin-driven iron competition further suggests that modulating iron availability could reduce *E. ruminantium* replication. Future research integrating metagenomics with transcriptomics and proteomics will be essential to disentangle microbiome-pathogen interactions and develop targeted interventions for reducing vector competence and heartwater-driven livestock losses across sub-Saharan Africa.

Several methodological considerations should be acknowledged. The sample size (11 infected/uninfected ticks per tissue) from a single location provides sufficient power for exploratory microbiome analysis but limits broader generalisability. That said, tissue-specific microbiome restructuring patterns broadly consistent with our findings have recently been reported in dromedary camel ticks (Khogali et al., in press), suggesting that pathogen-driven community reorganisation may be a generalisable feature of ixodid microbiomes rather than specific to *Am. gemma*. PICRUSt2 functional predictions serve as valuable hypothesis-generating tools but require experimental validation. The cross-sectional design prevents causal inference regarding microbiome changes. Finally, detection of *Rickettsia* reads in some pathogen-free samples likely reflects endosymbiotic rickettsiae commonly found in ixodid ticks (14, 33), highlighting the complexity of tick-associated microbial communities that merits further investigation.

Specifically, if *E. ruminantium* directly drives the observed shifts, controlled infection of pathogen-free *Am. gemma* colonies should reproduce *Luteimonas* expansion and *Rickettsia* depletion within the midgut over a defined post-infection window. Experimental infection models and integrated multi-omics approaches will be essential for moving from association to mechanism, and ultimately for developing targeted microbiome-based strategies to reduce the burden of heartwater on livestock production across sub-Saharan Africa.

## Acknowledgements

We are grateful to Dr. Daniel Masiga and Dr. Joel L. Bargul for their supervision and guidance throughout this study. We also extend our sincere appreciation to Epaphrus Yuko and David Wainaina of *icipe* for their invaluable assistance during fieldwork, and Jaqueline Wahura Waweru of icipe’s Bioinformatics Unit for her help with the initial data analysis. We further thank the farmers and veterinary staff of Kajiado County for their generous support and cooperation during tick collection from animals.

## Funding

This project has received funding from the European Union’s Horizon 2020 research and innovation programme under grant agreement no. 101000365 (PREPARE4VBD), and *icipe* institutional support from the Swedish International Development Cooperation Agency (Sida); the Swiss Agency for Development and Cooperation (SDC); the Australian Centre for International Agricultural Research (ACIAR); the Government of Norway; the German Federal Ministry for Economic Cooperation and Development (BMZ); and the Government of the Republic of Kenya. D.C. received training support through the Fogarty International Center of the U.S. National Institutes of Health, Eneza Data Science programme (Award 1UE5TW012539). The views expressed herein do not necessarily reflect the official opinion of the donors.

## Ethical approval and permits

The study was approved by the Pwani University ISERC (Ref: ERC/EXT/002/2020E) and licensed by NACOSTI (Ref: NACOSTI/P/24/38943). Verbal informed consent was obtained from livestock owners and documented in line with the approved procedures, as written consent was not feasible given variable literacy levels among participating farmers.

## Conflict of interests

The authors declare no conflicts of interests.

## Author Contributions

D.G., S.M., and J.V.: conceptualization. J.V.: project administration, and funding acquisition.

S.M. and J.V.: supervision. D.G., J.K., and R.K.: fieldwork and investigation. D.G., D.C., and J.V.: formal analysis and data curation, D.G.: writing – original draft. All authors reviewed and approved the final manuscript.

## Data, metadata, and code availability

All data, metadata, and code associated with this study are publicly available. Raw Oxford Nanopore Technology (ONT) reads, together with associated metadata, were deposited at the NCBI Sequence Read Archive (SRA) under BioProject accession number PRJNA1422828.

## Supplementary Information

**Figure S1**. Rarefaction curves showing observed bacterial OTUs as a function of sequencing depth in tick tissue samples. Each line represents an individual sample.

**Table S1.** ONT read tracking summary across bioinformatic processing stages.

**Table S2.** Top hub genera identified in tissue-specific microbial co-occurrence networks

**Table S3**. FDR-significant enzyme commission (EC) numbers differentially abundant between *Ehrlichia ruminantium*-infected and uninfected *Amblyomma gemma* microbiota.

**Table S4**. FDR-significant KEGG Orthology (KO) functions enriched in *Ehrlichia ruminantium*-infected *Amblyomma gemma* microbiota

